# Live tracking of a plant pathogen outbreak reveals rapid and successive, multidecade episome reduction

**DOI:** 10.1101/2023.05.23.541994

**Authors:** Veronica Roman-Reyna, Anuj Sharma, Hannah Toth, Zachary Konkel, Nicolle Omiotek, Shashanka Murthy, Seth Faith, Jason Slot, Francesca Peduto Hand, Erica Goss, Jonathan M. Jacobs

## Abstract

Quickly understanding the genomic changes that lead to pathogen emergence is necessary to launch mitigation efforts and reduce harm. Often the evolutionary events that result in an epidemic typically remain elusive long after an outbreak, which is particularly true for plant pathogens. To rapidly define the consequential evolutionary events result in pathogen emergence, we tracked in real-time a 2022 bacterial plant disease outbreak in US geranium (*Pelargonium* x *hortorum*) caused by Xhp2022, a novel lineage of *Xanthomonas hortorum*. Genomes from 31 Xhp2022 isolates from seven states showed limited chromosomal variation, and all contained a single plasmid (p93). Time tree and SNP whole genome analysis estimated that Xhp2022 emerged in the early 2020s. Phylogenomic analysis determined that p93 resulted from cointegration of three plasmids (p31, p45, and p66) present in a 2012 outbreak. p31, p45 and p66 were individually found in varying abundance across *X. hortorum* isolates from historical outbreaks dating to 1974 suggesting these plasmids were maintained in the broader metapopulation. p93 specifically arose from two co-integration events from homologous and Tn*3* and XerC-mediated site-specific recombination. Although p93 suffered a 49kb nucleotide reduction, it maintained critical fitness gene functions encoding, for example, metal resistance and virulence factors, which were likely selected by the ornamental production system. Overall we demonstrate how rapid sequencing of current and historical isolates track the evolutionary history of an emerging, ongoing threat. We show a recent, tractable event of genome reduction for niche adaptation typically observed over millenia in obligate and fastidious pathogens.

**Significance:** Genome-resolved epidemiology is rapidly changing how we track pathogens in real-time to support stakeholders and health. This research highlights how we responded to a current disease outbreak of geranium. Our work revealed that a new group of the bacterial plant pathogen *Xanthomonas horotrum* emerged in 2022 as a result of a recent genome reduction. We determined that three distinct plasmids were present in the broader *X. hortorum* metapopulation since 1974. In 2012, the three plasmids were altogether present in individual isolates; then in 2022, all three plasmids co-integrated while maintaining critical fitness genes but losing extraneous genomic material. This parallels genome efficiency and reduction that we see across millenia or even millions of years with obligate parasites with increased niche-specificity.

## Introduction

Emerging pathogenic microorganisms threaten human, animal, and plant health. Pathogenic bacteria adapt to changing environments through gene mutations and mobilization. Gene movement occurs within bacterial genomes and between cells. Intracellular DNA movement is usually mediated by Insertion Sequences (IS), transposases (Tn), and homologous recombination, and among cells, intercellular movement is caused by Horizontal gene transfer (HGT). HGT mechanisms are transformation (uptake of extracellular DNA), conjugation (plasmids), and transduction (mediated by bacteriophages).

Several disease outbreaks and their global dissemination are associated with plasmid-coded genes (1, 2). Plasmids are functional genetic modules that mediate gene transfer, conferring clinically and economically important properties to bacteria (1–5). An example of bacterial adaptation is the global dissemination of antimicrobial resistance genes threatening public health (6). The constant use of antibiotics has led to an increase in hospitals of Methicillin-resistance *Staphylococcus aureus* (MRSA). Another example is plant-associated bacteria with copper-resistance genes. Copper is a broad-spectrum biocide used to control foliar pathogens. Frequent application of copper led to the emergence of *Xanthomonas* and *Pseudomonas* copper-resistant isolates (7). Global outbreaks of plant pathogenic bacteria can serve as models to define pathogen emergence mechanisms.

A long-term evolutionary impact of parasitism is genome reduction (8). Over millions of years, hyper niche-specific microorganisms across the symbiotic spectrum experience major genome loss and often are obligate to their host (9). These million-year-long genomic changes are hard to track in real time but play fundamental roles in host adaptation and pathogen specialization (10, 11). Examples of genome reduction include bacterial pathogens, such as the 1.1Mb genome of *Rickettsia sp*. that causes typhus fever, the *Burkholderia mallei* chromosome size reduction that causes equine disease, and the Xanthomonadacea xylem-limited plant pathogenic bacteria *Xylella fastidiosa* (2Mb genome) and *Xanthomonas albilineans* (3Mb genome) causing Olive Quick decline and leaf scald in sugarcane (10–12).

The environmental horticulture industry, including floriculture crops, is core to the US economy, totaling $4.80 billion. Among these, geranium plants are particularly important, valued at $5.9 million in 2020 (USDA ERS). A destructive disease of geraniums, reported worldwide, is bacterial blight caused by *Xanthomonas hortorum* pv. pelargonii (Xhp) (13). Xhp spreads by plant vegetative- and seed-propagation, irrigation, and wounds (14–16). The reported symptoms in the leaves are water-soaked spots that turn black, wilting leaf margins, or chlorotic V-shaped lesions (Figure 1A). Stems develop black rot and the whole plant wilts. Several Xhp outbreaks have been reported with 10-100% estimated annual losses in greenhouses and in-field conditions (17–20). In 2022, nationwide producers reported a foliar outbreak of geranium bacterial blight caused by *Xanthomonas hortorum* pv. pelargonii (21). Even with recent reports tracking Xhp, little is known about Xhp evolution, functional diversification, and pathogen epidemiology (22).

**Figure 1.**
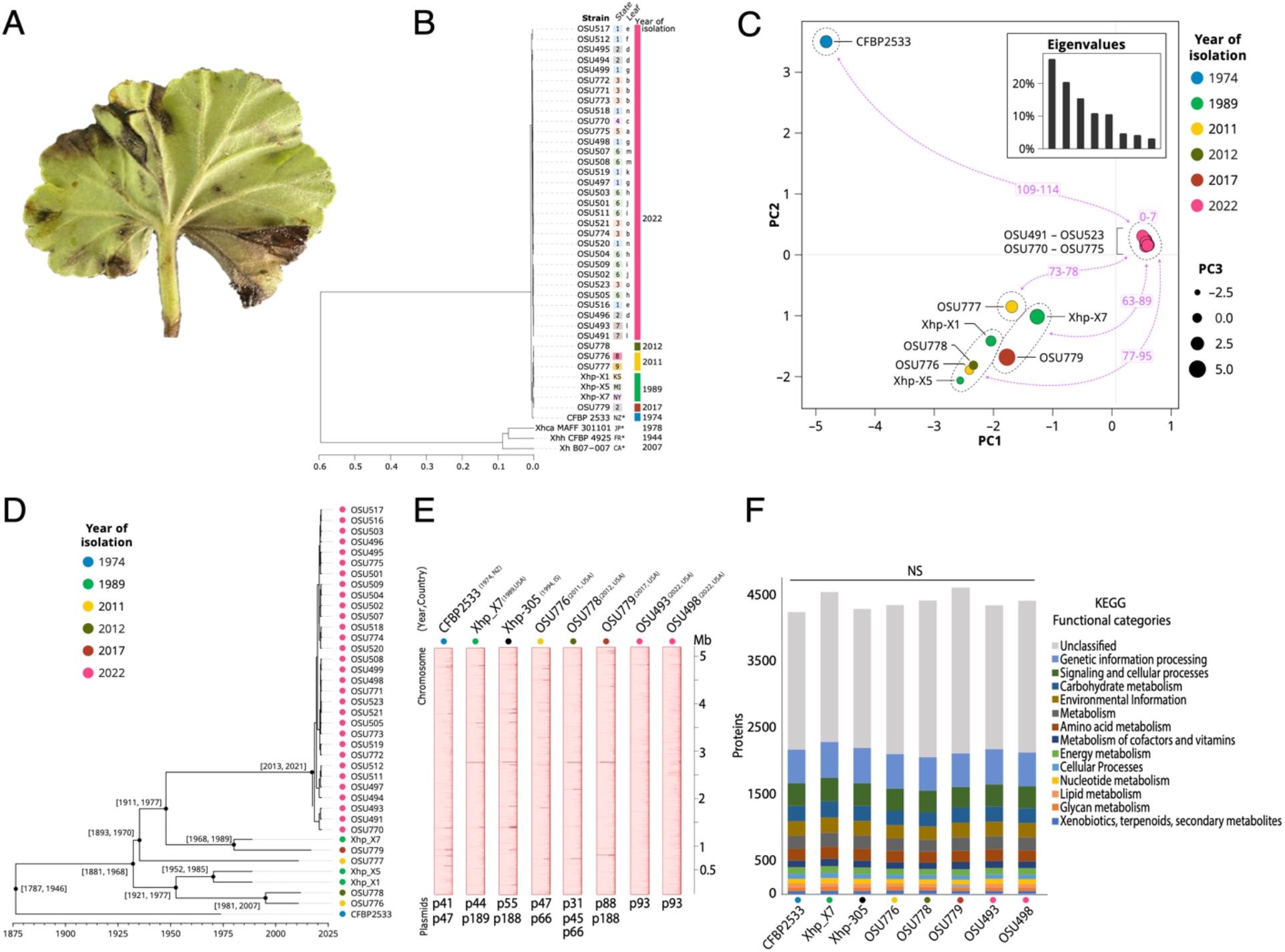
Genomic identification and characterization of the recent geranium Xh2022 bacterial blight outbreak. A) Symptoms of blight in a geranium leaf. B) Whole genome Average Nucleotide Identity analysis. Numbers in colored squares indicate geographic location. Lowercase letters show isolates from the same leaf. Colored rectangles indicate the collection year. Xhp, *X. hortorum* pv. pelargonii. Xhh, *X. hortorum* pv. hederae; Xhca, *X. hortorum* pv. carotae) Clustering of the Xhp isolates based on principal component analysis of core chromosomal SNPs. The number of SNPs differentiating the cluster of 2022 isolates from other clusters are shown in purple. D) Time-scaled phylogeny of Xhp using an HKY nucleotide substitution model. Brackets indicate 95% HPD credible interval for date of each node. Colored dots indicate the collection year. E) Mauve Multiple Xhp Chromosome alignment. Each chromosome is rotated to start with the DnaA sequence. Each color shows genetically similar blocks. F) KEGG functional annotation of Xhp chromosomes.

Advances in genome sequencing applications for genome-resolved epidemiology have allowed immediate tracking of the evolutionary history of human clinical outbreaks but remain to be widely employed for plant disease epidemics (23, 24). Rapid sequencing and public data dissemination have made the surveillance process an international effort to mitigate economic and food security risks (25). In 2022, there was an outbreak of geranium blight caused by *X. hortorum* pv. pelargonii. We, therefore, quickly (two days) used whole genomes from infected leaf tissue to define the nature of the evolutionary of the geranium outbreak.

## Results

### The 2022 geranium bacterial blight outbreak is due to the emergence of a novel Xhp lineage

We received 31 Xhp isolates from infected geranium collected in 2022 from 18 different USA greenhouses across seven states (Table S1). To define the pathogen’s identity and evolutionary history of the 2022 nationwide outbreak compared to historical events, we used short-read Illumina sequencing to generate genome assemblies for isolates from the current and historical outbreaks: 31 from 2022 (Xhp2022); three from 1989 (X-1, X-5, X-7), two from 2011 (OSU776, OSU777), one from 2012 (OSU778) and one from 2017 (OSU779) (21). For the analysis, we used the Xhp type-strain CFBP2533 from 1974 as a reference (26). CFBP2533 was the only Xhp genome available on NCBI when we started the analysis in 2022.

To examine genomic distances among the 38 sequenced Xhp isolates, we calculated average nucleotide identity (ANI) among all isolates and included three non-geranium pathogen *X. hortorum* as outgroups. ANI comparisons showed all 38 isolates belonged to the pathovar (pv.) pelargonii and had 96% ANI compared to *X. hortorum* pv. hederae NCPPB939, *X. hortorum* pv. carotae MAFF301101 and *X. hortorum* B07-007 (Fig 1B). Within the 38 Xhp, Xhp2022 isolates formed a single cluster (99.95% ANI) with no location-related clusters (Fig. 1B).

We then identified core-genome Single Nucleotide Polymorphism (SNP) for all Xhp isolates (Fig. 1C). The principal component analysis showed five groups based on SNP genotypes. The Xhp2022 isolates formed a distinct group with zero to seven SNPs among them, confirming the ANI comparisons. All other isolates were differentiated from the 2022 outbreak by 63 to 114 SNPs.

To determine when the new Xhp2022 group diverged from other Xhp, we inferred a time-calibrated phylogenetic tree. The analysis revealed that Xhp2022 isolates emerged from a common ancestor dated to 2017 (95% highest posterior density interval: 2013-2021; Fig. 1D). All the genomics analyses showed that the recent outbreak is due to the emergence of a novel lineage.

### Older and recent Xhp isolates have similar chromosome content and structure

To further define the genome structure including chromosome and plasmid resolution in Xhp2022 compared to previous outbreaks, we generated long-read, Oxford Nanopore complete genomes of seven Xhp isolates from different years: 1989 (X-7), 2011 (OSU776), 2012 (OSU778), 2017 (OSU779), and 2022 (OSU493, OSU498) (Table S1). We used Xhp CFP2533 from 1974 as a reference, as the complete genome is publicly available on NCBI (27). We included long-read sequenced strain Xhp-305 isolated in 1994, as it was uploaded to NCBI in February 2023. Based on the assemblies, there were few differences in whole chromosome synteny across six decades of Xhp isolates. Instead, there was a high variation in plasmid abundance and sizes (Fig. 1E). Based on the assemblies, we compared chromosomes and plasmids separately.

The Xhp chromosomes alignment showed no rearrangements or inversions and one contiguous colinear block (Fig. 1E). The total chromosomal gene content was 4455±200 (Fig. 1F, Table S2) with no significant differences across isolates (F (7,104) = 0.01, p=0.99). The most abundant genes encoded mobile genetic elements (MGE), which are associated with gene movement and host adaptation (Table S2) (28). Therefore, we annotated the chromosomes using the mobile genetic elements database mobileOG-db (29). Overall, all seven chromosomes have 202 to 221 MGEs, and shared most of their locations (Fig. S1A, Table S3). Notably, these genomes over a 60-year time span did not show significant differences in quantity and location (F (7,32) = 0.011, p=0.99).

Pathogenicity factors such as Type three secreted effectors (T3E) are injected directly into plant cells to manipulate immunity and required for successful host interaction (30). To assess the T3E diversity across Xhp chromosomes, we annotated the chromosomes for Type three secreted *Xanthomonas* Xop effectors. Chromosomes from all seven long-read genomes have 19 to 20 Xop effectors and shared 14 of them in the same loci. (Fig S1B, Table S4).

Next, we assessed codon usage differences across Xhp isolates to determine if the chromosome is under pressure to increase translation efficiency for bacterial fitness (31). Based on Dynamic Codon Biaser (DCB) tool, all Xhp chromosomes have a similar codon usage to Xhp CFBP2533 (R^2^=0.999) (Table S8). Overall, location and content of host-associated fitness factors and MGEs indicate the Xhp chromosomes were similar across decades.

### The Xhp 2022 isolates have a reduced episome

Plasmids are a primary vector for bacterial horizontal gene transfer (HGT) and rapid adaptation to environmental change (4). The long-read assemblies revealed that Xhp isolates from previous outbreaks have at least two plasmids, from 31Kb to 189Kb. The Xhp2022 isolates only encoded one plasmid (p93) of 93Kb. Based on plasmid number and size differences among Xhp isolates, we explored the plasmid structure and gene content.

To determine the similarity in p93 plasmid content across Xhp2022 and other *Xanthomonas* species, including historically Xhp sequenced above, we determined regions of plasmids with shared identity with p93. BLASTn searches revealed that Xhp2022 p93 contained three distinct regions with variable functions (Fig. 2A, Table S5). Region 1 had high nucleotide identity (99%) and 20% coverage with Xhp isolates from 1989 (p189), 1994 (p188), 2011 (p66), 2012 (p66), and 2017 (p189) (Fig 2A). *X. citri* pv.citri and *X. euvesicatoria* plasmids, and *Stenotrophomonas maltophilia* chromosome aligned (92-93% identity) to Region 1, which highlights the conservation and mobility of this region outside Xhp. Region 2 was also present in plasmids across Xhp isolates with 99% identity (30% coverage) from 1974 (p47), 1989 (p44), 1994 (p55), 2011 (p47), 2012 (p31), 2017 (p88), and *X. hortorum* Oregano_108 plasmid2 (Fig. 2B), followed by 95-97% identity to other *X. hortorum* pathovars, and *X. citri* pathovars fuscans and phaseoli. Region 3 shared identity (97-99%, with 50% coverage) to the plasmids from 1974 (p41), 2012 (p45), *X. campestris* pv. *campestris*, other *X. hortorum* pathovars and *X. citri* pathovars (Fig. 2C). Additional BLASTn using each 2012 plasmid as query confirmed the three Xhp2022 p93 regions with 99% identity and with more than 60% coverage (Table S5). A search in the plasmid database PLSDB for p93 and the 2012 plasmids validated the BLASTn results (32).

**Figure 2.**
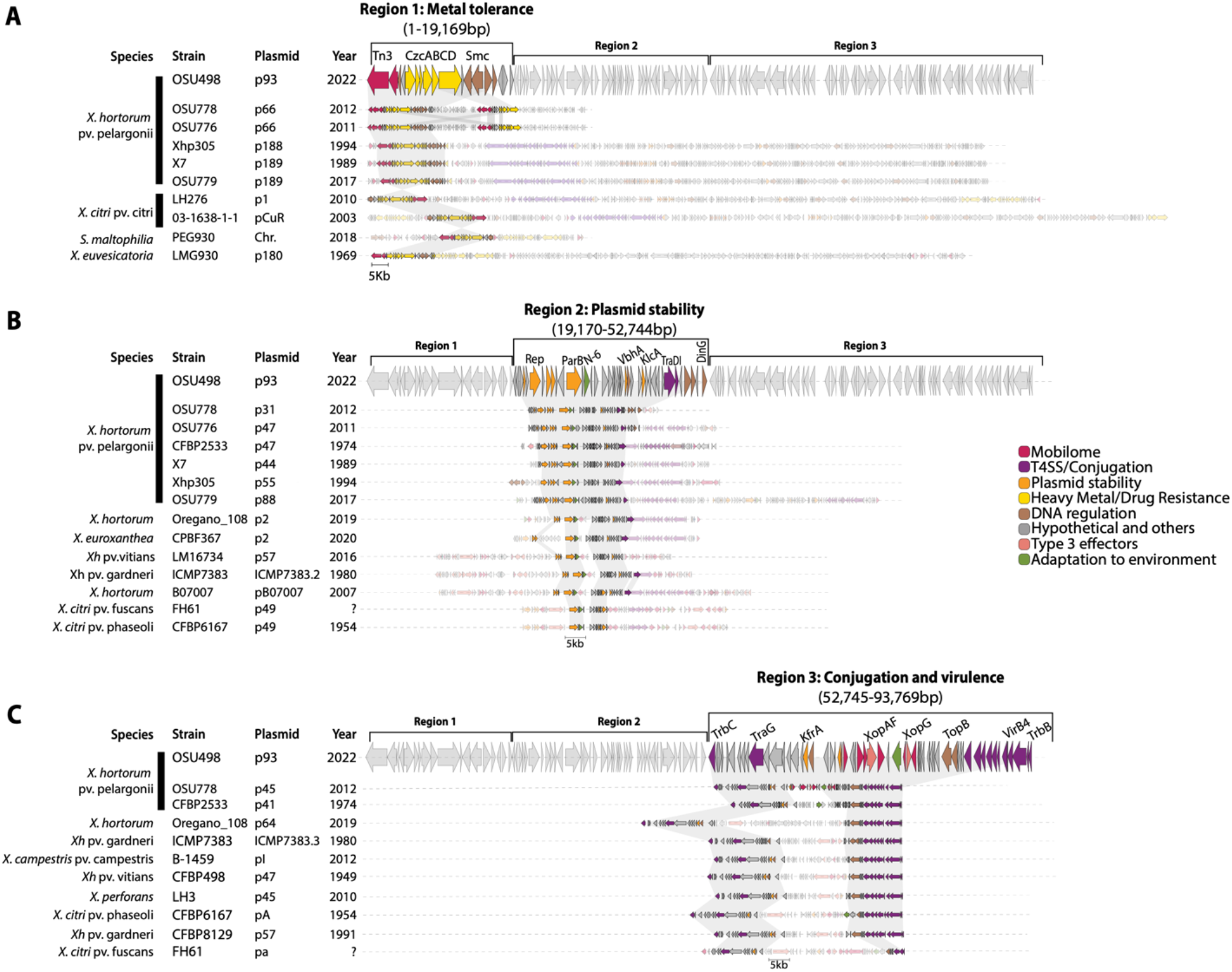
Gene cluster alignments of Xhp2022 and *Xanthomonas* plasmids. Clinker plasmids protein alignments to the A) Region 1: Metal Tolerance B) Region 2: Plasmid Stability and C) Region 3: Conjugation and Virulence Strategies of p93. Each arrow represents a coded gene and color represent a broad classification. The gray translucent connecting squares indicate similarities among plasmids that are above 60%.

For gene coding sequence comparisons, we performed a global clinker alignment for each p93 region with at least nine different *Xanthomonas* species plasmids (23). The alignment showed the p93 regions had similar structure and protein content to the other nine plasmids (greater than 60% alignment sequence similarity) (Fig. 2). Region 1 contained proteins associated with cobalt/zinc/cadmium efflux system CzcABCD, mobile elements (Tn3 and recombinases), structural maintenance of chromosomes (Smc) and DNA binding (SEC-C, and exoribonuclease). Region 1 shared proteins with the largest Xhp plasmids from 1989, 1994, 2011, 2012, and 2017. All Xhp plasmids shared the same CzcABCD cluster with *X. citri, X. euvesicatoria* pLMG930.1, and *S. maltophilia* chromosome (Fig. 2A). Region 2 had genes coding for DNA replication and regulation proteins (Rep, ParB, 3’-5’ exonuclease DinG, and RecD), bacterial fitness (N6 DNA Methylase), and antitoxins (VbhA, RelE/ParE, VapC). The Region 2 shared these proteins with Xhp plasmids from 1974, 1989, 1994, 2011, 2012, and 2017. These proteins are also present in five different *X. hortorum* isolates, *X. euroxanthea* and two different *X. citri* pathovar plasmids (Fig. 2B). Region 3 had genes coding for proteins associated with conjugation (Vir and Tra protein cluster), T3E (XopAF and XopG), and DNA replication (DNA topoisomerase TopB). Region 3 shared all proteins with the plasmid p45 from 2012 and partly with p41 from 1974. Additionally, it shares most of the content with other *X. hortorum* pathovars, *X. citri* pv. citri, *X. perforans* and *X. campestris* (Fig. 2C). Overall, the p93 regions appear to have conserved functions among older Xhp plasmids and other *Xanthomonas* species plasmids.

### The Xhp2022 episome emerged by plasmid reduction by homologous recombination

Next, we annotated the plasmids using mobileOG-db to assess the plasmid mechanisms for gene movement and stability (29). The three Xhp2022 p93 regions overlapped with p66 and p45 at genes coding for Integration/excision proteins (Tn3, xerC, recombinase, invertase) and with p31 and p45 at genes encoding DNA repair enzymes (dinG, recBCDF, Exoribonuclease), conjugation machinery proteins (trb genes), and with p66 and p31 at genes encoding DNA binding enzymes (smc, sec-C) (Fig. 3A). The overlapping genes suggested homologous and site-specific recombination as potential mechanism for plasmid cointegration (Fig. 3B) (2, 33).

**Figure 3.**
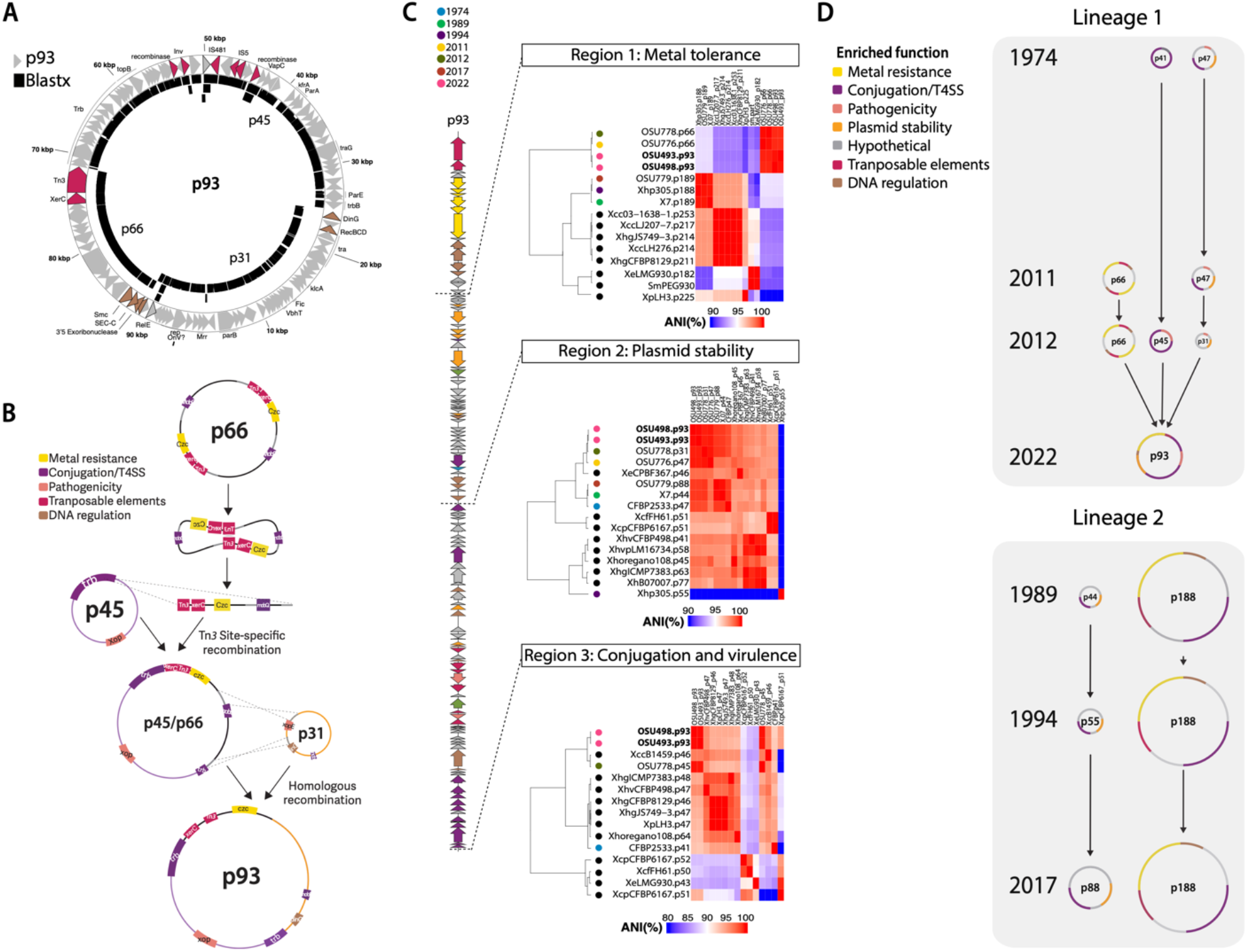
Evolutionary dynamics of plasmid co-integration in Xhp2022. A) Overlapping genes among 2022 and 2012 plasmids. The arrows represent p93 genes and the black blocks represent BLASTx (identity> 90%) analysis of each 2012 plasmid against p93. Pink arrows represent transposable elements and brown arrows represent DNA regulation-associated genes. B) representation of the cointegration events that led to the p93. First a Tn3-xerC site-specific recombination event integrated p66 into p45. The identical regions between p31 and p66 allowed a homologous recombination event that integrated p31 into p93. C) Average nucleotide identity comparisons heatmaps among p93 regions, Xhp plasmids and other *Xanthomonas* plasmids. The colored circles represent the isolation years for each Xhp isolate. Black circles are other *Xanthomonas* plasmids from NCBI. D) Xhp plasmid evolution model. Translucent gray squares indicate the two Xhp lineages based on the plasmid structures. Black lines represent the evolution path for each plasmid. Each color in the plasmids indicate enriched functions transfers across years.

To precisely find the expansion events in the Xhp2022 plasmid, we predicted the plasmid mobility, analyzed the phylogenetic relationships of individual proteins in each p93 region, and performed average nucleotide comparisons of whole plasmids. For plasmid mobility, we classified the Xhp plasmids by relaxase type (*mob*), and plasmid transferability (34). We found four relaxases, *mob*F, *mob*P, *mob*H, and *mob*Q. p93 had *mob*F and *mob*P (Table S6). Plasmids similar to Region 1 had either *mob*H or *mob*Q. The plasmids that matched Region 2 had *mob*F, and Region 3 had *mob*P. All plasmids were conjugative or mobilizable except for p44, p47, and p31 from 1989, 2011, and 2012 respectively. Loss of mobilization mechanisms may explain the driving force for plasmid co-integration and reduction.

To test the evolutionary origin of each region, we created phylogenetic trees based on representative proteins encoded in each region. From Region 1 we selected five genes encoding the Czc cluster and Sec-C. The Czc system is involved in heavy metal resistance (2, 35), and Sec-C is associated with protein secretion (36). The proteins CzcACD clustered with p66 from 2012 and other *X. hortorum* (bootstrap>99%) (Fig. S2A). CzcB and Sec-C associations were inconclusive due to bootstrap values lower than 90%. The protein clusters suggested that most genes from Region 1 come from Xhp but originated from *Stenotrophomonas* chromosome (bootstrap >98%) and probably then moved to other *Xanthomonas* species plasmids. For Region 2 we selected Rep, ParB, N6 DNA Methylase, VbhA, and DinG proteins. Rep, ParB, and N-6, clustered with *X. hortorum*, VbhA clustered with *X. hortorum* and *X. arboricola* (bootstrap > 98%) (Fig. S2B). The protein DinG clustered with several *Xanthomonas* species. The outgroups (bootstrap > 90%) suggested that proteins from Region 2 came from Xhp and likely have *Xanthomonas* species plasmids as common donors. For Region 3 we selected TadA, TraG, XopAF, XopG, and TopB proteins. All proteins grouped with *X. hortorum* but only TraG had a bootstrap value of 99% (Fig. S2C). The trees suggested that proteins from Region 3 originate directly from Xhp.

To estimate relatedness among Xhp plasmids in each region we calculated ANI values (Fig 3B). Plasmids associated with Region 1 formed two groups, the long plasmids from 1989, 1994, 2017, and plasmids from 2011, 2012, 2022. Region 2 and 3 indicated that these plasmids are common for *X. hortorum* isolates, and they formed a single cluster. Overall, our results show the origin for each region resulted from the 2012 outbreak with ancestral origins outside Xhp.

Next, we annotated the Type three effectors (T3E) as virulence and transmission traits for the pathogen (30). We found four effectors in Xhp plasmids: TAL effector, *xopAF2, xopG1*, and *xopE2* (Table S7). The TAL effector was annotated as a pseudogene and was only present in p47 from 1974. *xopAF2* and *xopG1* were present in p93 and p45 from 2012. *xopE2* was present in 1974, 2011 and 2012 plasmids but not in p93. Based on these results, the only virulence factors present in p93 came from p45 from 2012.

## Discussion

Microorganisms with strong niche-adaption display major genomic reduction (37). Chromosome reduction is an indicator of specialization in obligately host-associated organisms. To our knowledge, the bacterial extrachromosomal reduction has not been fully studied in plant pathogens. In this study, we suggest that an Xhp population became predominant due to plasmid reduction, which led to the bacterial geranium blight outbreak in 2022. The Xhp2022 plasmid (p93) has elements associated with gene movement and fitness that pointed us to propose that p93 originated from cointegration plasmid. We propose site-specific and homologous recombination as the events that led to the plasmid reduction (Fig 3B). Based on the structure of p66, the Tn*3*-*xerC* integrated within the *trb* operon from p45, as Tn*3* movement does not require similar sequences in the recipient plasmid. The integration in the *trb* operon could explain why *trbC* and *trbB* are no longer syntenic in p93. Since p66 is a dimer plasmid, it is possible that *xerC* resolved the dimer and only a monomer integrated into p45. The integrated p66 fragment has identical regions to p31 that could have led to homologous recombination to integrate p31. The recombination could have excised 20 genes from p66 among *mobQ*, and three genes from p31 (*xopE*, DNA-invertase *hin*, hypothetical protein). We suggest that the non-conjugative plasmids p66 and p31 cointegrated into p45 for mobilization, as cointegration is a mechanism for moving non-transferable plasmids from the same host (5, 38–40).

Based on plasmid structure and ANI comparisons of all the Xhp isolates, we propose two Xhp plasmid lineages (Fig 3C). Xhp isolates from 1974, 2011, 2012, and 2022 belong to Lineage 1, and isolates from 1989, 1994, and 2017, are part of Lineage 2. The lineages differences are the *trb* and *tra* locus, T3E, and two more operons associated with copper and arsenic tolerance (Fig 2). Likely, after plasmid cointegration, the Xhp population from Lineage 1 was enriched, which led to the 2022 outbreak. Plasmid diversification indicates bacterial adaptation to different environmental pressures, as has been described for other plant pathogens like the oncogenic plasmids from *Agrobacterium* (41).

Plasmid protein phylogenies showed that most genes in p93 are present in *X. hortorum* subgroups but originated from other *Xanthomonas* species or *Stenotrophomonas*. Moreover, Xanthomonadaceae species have higher HGT plasmid efficiency (42, 43). We suggest that several *Xanthomonas* species have similar selection pressures to promote the proliferation of plasmid-containing bacteria and genetic interchange across species (44, 45). That could explain why several plant pathogenic *Xanthomonas* species from different years, hosts, and continents have similar plasmids to p93 (identity >50%) (Fig S3). An example of shared selection pressure for Xanthomonadaceae is copper usage for disease management (46). Copper usage promotes the movement of the copper-tolerance genes from *X. citri* plasmids and *Stenotrophomonas maltophilia* chromosomes and vice-versa (35, 47).

*Xanthomonas* species have similar agricultural settings contributing to pathogen spread, like host and soil microbiome and greenhouse practices. The microbiome is a source of genetic interchange (48, 49). The soil and host microbiome could be a reservoir for copper-tolerance and virulence-related genes horizontally acquired by new isolates (45, 50, 51). Greenhouse facilities might have multiple crops, use vegetative propagation, and irrigation water (splashing water) that leads to pathogens spreading. These agricultural settings shared among plant pathogenic *Xanthomonas* might allow them to interact and share genetic materials for fitness.

Overall, we used short and long sequencing to describe the genomic structure of the 2022 Xhp outbreak and compare it to isolates from earlier decades. We provide a comprehensive description of the Xhp outbreak with single nucleotide polymorphisms and whole genome structure. We propose that an Xhp population with reduced plasmid become predominant leading to the 2022 outbreak. Based on the minimal differences in phylogenetic distances and SNP variations, we hypothesize that the Xhp2022 nationwide outbreak initial inoculum arrived from a single geranium production location abroad and was then disseminated across the United States, consistent with previous findings (22).

## Materials and methods

### Bacteria isolates and genome sequencing

*X. hortorum* pv. pelargonii isolates were obtained from US nurseries. Bacterial DNA extraction for short-read and long-read sequencing was done with the Monarch NEB kit and Qiagen kit, respectively. The library for short read sequences were prepared and sequenced at the Applied Microbiology Services Lab (AMSL)-Ohio State University. Briefly, Sequencing libraries were prepared with Illumina DNA prep (Illumina(R), San Diego, CA), multiplexed with 10 nt dual unique indexes (IDT, Coralville, IA) and sequenced at a target depth of 100x in Illumina NextSeq2000. Long-read sequences were generated with the Oxford Nanopore MinION, Flowcell 9.4, and the Rapid ligation kit (RBK004). Raw data and assembled genomes are deposited in the NCBI GenBank with the project (PRJNA846756).

### Short read analysis

Core chromosomal single nucleotide polymorphisms were identified using the PROK-SNPTREE pipeline (52). Briefly, the quality of the raw reads was verified with FASTQC v0.11.7 (bioinformatics.babraham.ac.uk/projects/fastqc) and the adapters were trimmed with TRIM_GALORE v0.6.5 (53). Clean reads files were aligned to whole genome assembly of *X. hortorum* pv. pelargonii strain CFBP 2533 (27) using BWA v0.7.17 (54). Variants calling was performed using the HAPLOTYPECALLER tool in GATK V4.1.9.0 to generate base-pair resolved Variant Calling Format (VCF) files (55). For each SNP position, the nucleotide base for each isolate at that position was determined from base-pair resolved VCF file under stringent filtration parameters (absent if QD < 5; reference allele if Alternate allele Depth < 10 × Reference allele Depth; and else alternate allele) to generate an allele table. All chromosomal positions with high-quality SNPs were retained.

The nucleotides for positions conserved in all isolates were concatenated to generate SNP-alignment which was used for subsequent analyses. Principal component analysis was performed in R v4.2.2. A distance matrix was generated from SNP-alignment using the r/adegenet package (56) and the principal components were computed using r/ade4 package (57). The dated phylogenetic tree was generated from the SNP-alignment and year of collection in Bayesian Evolutionary Analyses of Sampled Trees (BEAST) v. 2.7.3 (58). Priors included, HKY nucleotide substitution model with empirical base frequencies, an uncorrelated relaxed clock, and Bayesian Skyline (59). Markov Chain Monte Carlo chain was run for 100 million generations, which produced minimum effective sample sizes of 1000 for each estimated parameter. From the posterior distribution of 10,001 trees, the maximum clade credibility tree was derived using TREEANNOTATOR v2.4.0 (60) and subsequently visualized in FIGTREE v1.4.4 (tree.bio.ed.ac.uk/software/figtree).

For whole genome analysis, raw reads were cleaned using Trimmomatic and de-novo assembled using Unicycler and Spades (61–63). Scripts are available in https://github.com/htoth99/Geranium_Project_2022/. Enveomics website was used for whole genome Average Nucleotide Identity comparisons and tree generation.

### Long read analyses

Long raw data was based-called and demultiplexed with Guppy v5 (https://community.nanoporetech.com). Samples were assembled using Flye v. 2.29.2, polished with Homopolish v0.4.1, and annotated using Prokka v1.14.5 (64–66). The chromosomes were rotated to the DnaA using the python code from Dr. Ralf Koebnik (IRD) and aligned using Progressive Mauve Alignment (67). Functional KEGG categories were assigned using BLAST Koala (68). The effectors for the chromosome and plasmids were predicted using a Xop *Xanthomonas* proteins database and BLASTx. To determine the Mobile genetic elements, we used the MobileOG-db program thru the website Proksee (69). To determine codon usage in the chromosomes we used the website program Dynamic Codon Biaser (www.cdb.info).

To determine the plasmid relaxases type and their transferability, we used the program MOB-Suite v3.1.2 and the tool MOB-typer (34). To identify similarities to other plasmids, we used the fasta file of each plasmid and used the website database PLSDB with “Mash dist” search strategy, and 0.1 as the maximum p-value and distance (32). To describe the plasmid similarities with the NCBI nucleotide database, we use the plasmids p93 and the 2012 plasmids as query in BLASTn. We kept the first ten hits with >52% coverage and >90% identity. Based on the BLAST results we selected 5-10 plasmids for protein comparisons. We downloaded the complete accession for each plasmid and annotated with Prokka to have the same annotations. We then perform a whole plasmid protein alignment using the program, Clinker (70). The results analysis with each plasmid was exported as a html and only connection with more than 60% similarity were kept. Colors were edited in Adobe Illustrator.

Based on NCBI and clicker results, we selected five proteins from each plasmid region for further analysis. We used the algorithm BLASTp, the protein as query, and limited the search for the group Xanthomonadaceae. The 100 hits for each gene were downloaded for phylogeny reconstruction. Proteins were aligned and clean using MAFFT and ClipKIT (71, 72). Then we constructed trees using the program FastTree to check the root (73). Based on the Fastree results we run the analysis using at least 50 sequences with IQ-TREE. Trees were visualized with Figtree. The trees were rooted based on the highest bootstrap value. Whole plasmid Average Nucleotide Identity analysis and trees were done with the program Pyani and the ANIb format (74).

Xanthomonadaceae whole genome tree was build using the website Enveomics and only using complete genomes (chromosome and plasmids) (75). Tree was plotted using the Ward model.

## Acknowledgements and funding sources

We are grateful for our funding sources including: USDA NIFA FACT-CIN (No. 2021-67021-34343, Ohio State’s Infectious Diseases Institute’s Interdisciplinary Seed Grant Program and the Ohio Department of Agriculture’s Specialty Crop Block Grant to JMJ. We thank the Ohio Super Computer for bioinformatics support, and Dana Martin and Taylor Klass for technical support for *Xanthomonas* isolation and sample processing.

---

**Figure S1.**
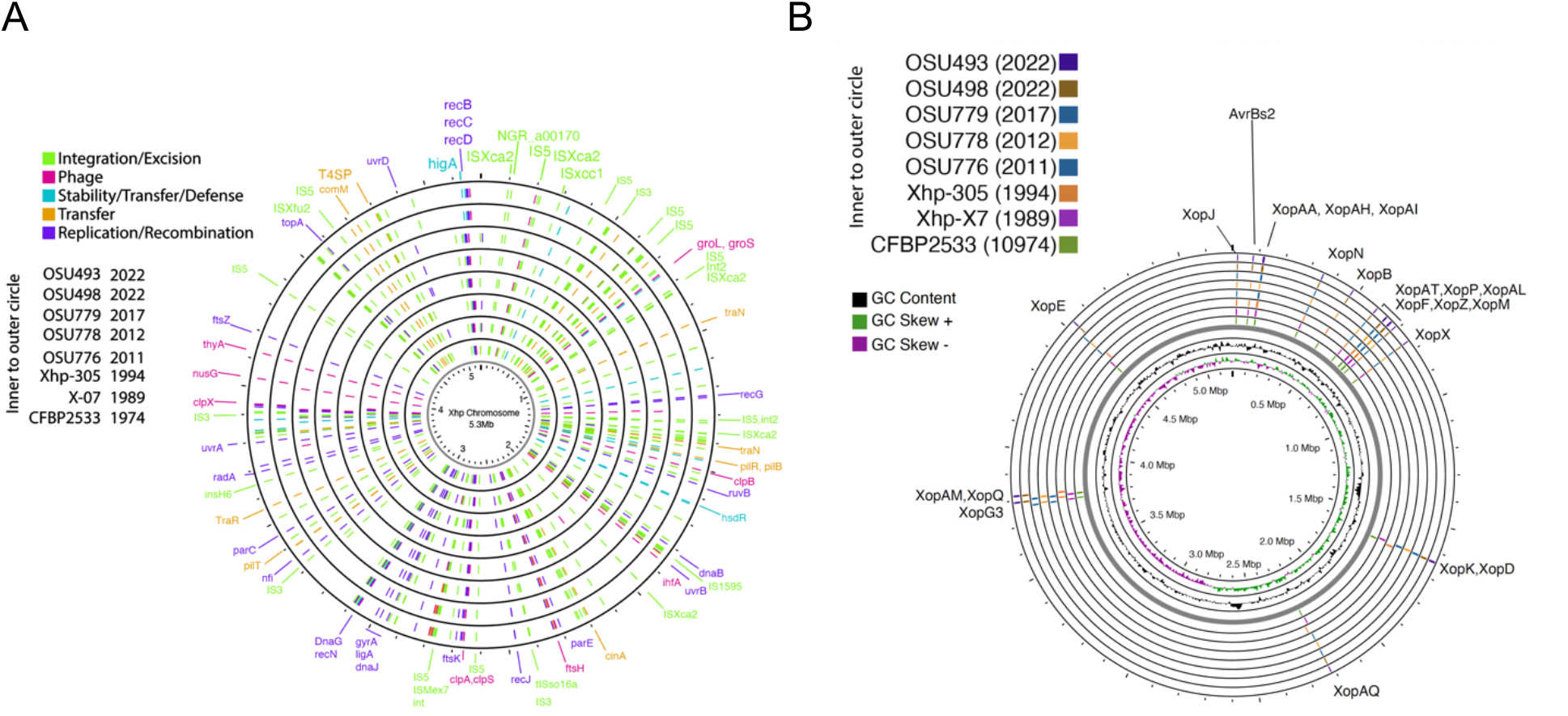
Mobile genetic elements and virulence factors in Xhp Chromosomes A) Mobile genetic elements in each Xhp chromosome annotated with MobileOG-db. B) Effector content in the Xhp chromosomes using BLASTX search. Effectors were considered present with a percentage of amino acid identity higher than 50% and coverage higher than 70%. Each ring represents a genome all aligned to dnaA sequence. Graphs and MGE annotations were made with Proksee (66).

**Figure S2.**
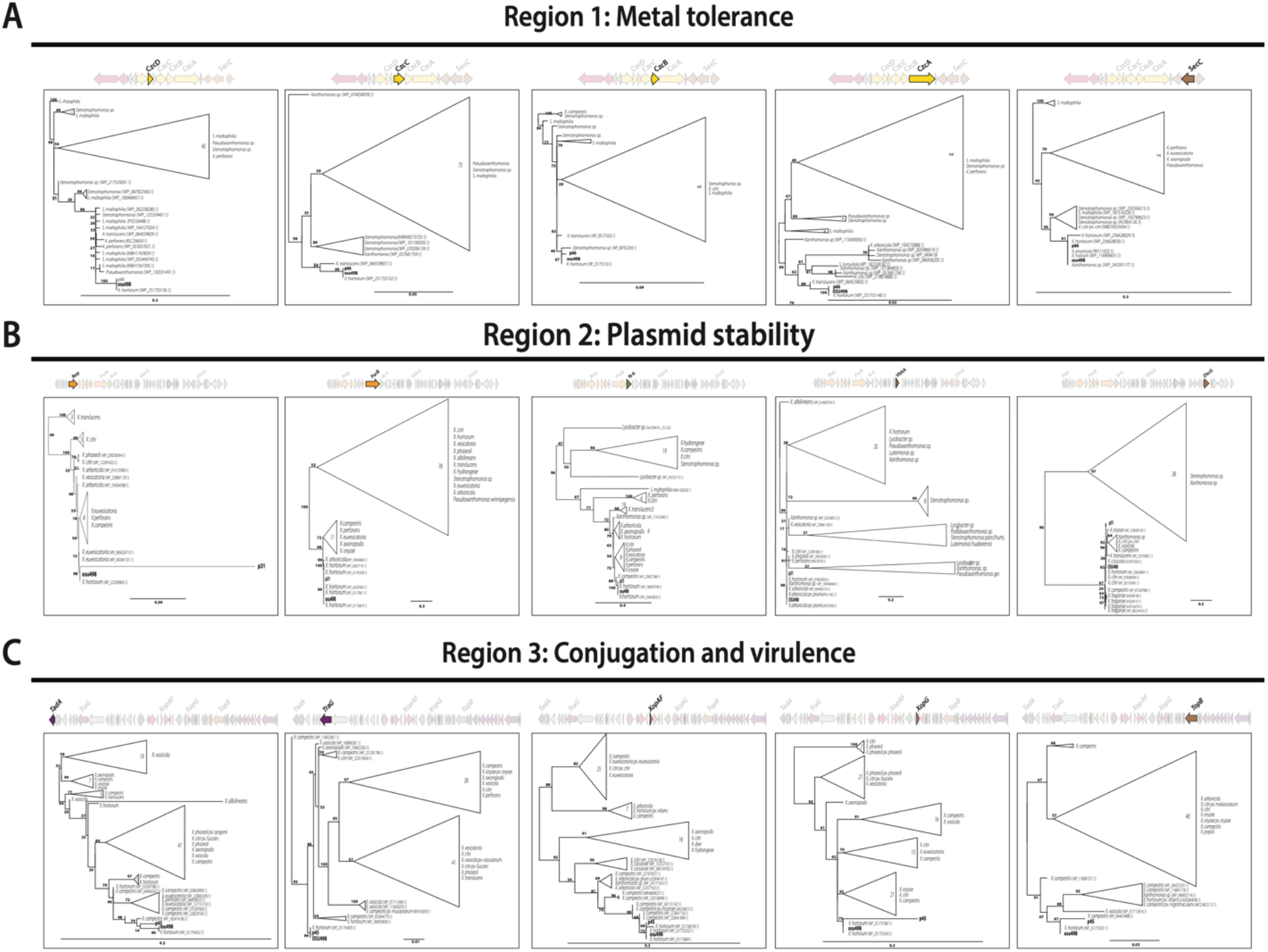
Phylogenetic comparisons of proteins coded in each Xhp plasmid shared among Xanthomonadaceae. Proteins were aligned using MAFFT. Alignments were trimmed with ClipKIT and trees were constructed with IQ-TREE. The outgroup for each tree was defined based on highly supported nodes within that tree (bootstrap>90) and a highly supported descendent node.

**Figure S3.**
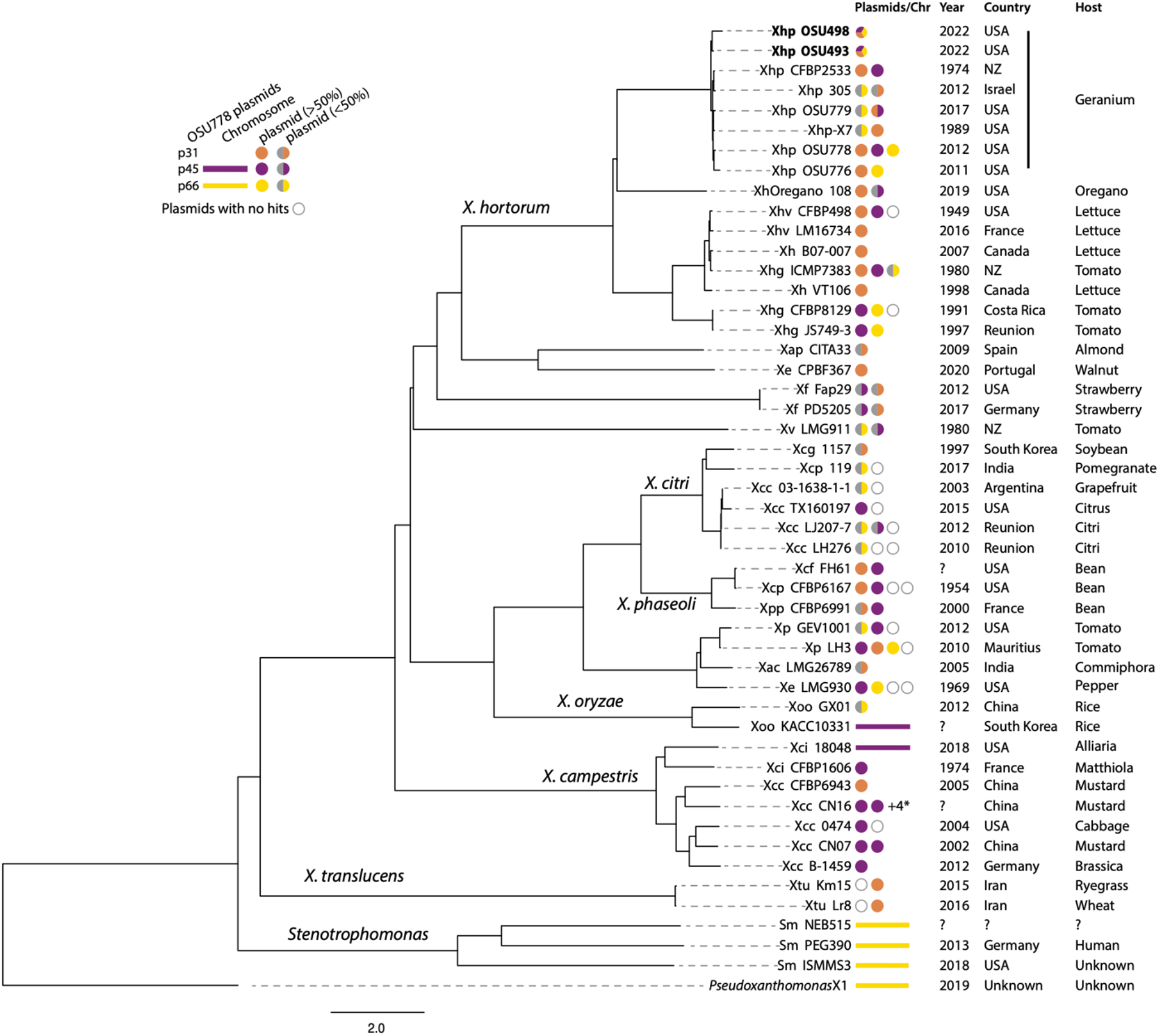
Phylogenetic tree of bacterial genomes with hits to Xhp 2012 plasmids. Yellow represents hits to p66, purple represent hits to p45, and orange represent hits to p31. If the plasmid shared 10-49% genes with the plasmid, half of the circle is gray; if the plasmids have more than 50% nucleotide identity, the circle has one color. The tree was build using Average Nucleotide Identity on the Enveomics website and visualized as a phylogenetic tree and the Ward model. +4* indicated plasmids on NCBI that had a size less than 5Kb. Question mark indicates no information in the NCBI Biosample.

## Notes

### Competing Interest Statement

The authors have declared no competing interest.

